# COGNITIVE REJUVENATION IN OLD RATS BY HIPPOCAMPAL OSKM GENE THERAPY

**DOI:** 10.1101/2023.06.13.544719

**Authors:** Steve Horvath, Ezequiel Lacunza, Martina Canatelli Mallat, Enrique L. Portiansky, Maria D. Gallardo, Robert T. Brooke, Priscila Chiavellini, Diana C. Pasquini, Mauricio Girard, Marianne Lehmann, Qi Yan, Ake T. Lu, Amin Haghani, Juozas Gordevicius, Martin Abba, Rodolfo G. Goya

**Affiliations:** Altos Labs, San Diego, USA; Human Genetics, David Geffen School of Medicine, University of California, Los Angeles, Los Angeles, California, USA; Epigenetic Clock Development Foundation, Torrance, California, USA; CINIBA, School of Medicine, UNLP, La Plata, Argentina; Institute for Biochemical Research (INIBIOLP) - Histology B & Pathology B, School of Medicine, National University of La Plata (UNLP), La Plata, Argentina; Institute of Pathology, School of Veterinary Siences; Vitality in Aging Research Group (VIA), Fort Lauderdale, FL, USA

**Keywords:** Hippocampal aging, spatial memory, OSKM gene therapy, rejuvenation, OSKM-induced demethylation, epigenetic age

## Abstract

Impaired performance in spatial learning and memory during aging in rats is associated with morphological and molecular changes in the brain, particularly in the hippocampus. Here, we assessed the cognitive performance of young (3.5 mo.) untreated rats and old (25.3 mo.) treated and control rats. Treatment was carried out by intrahippocampal injection of an adenovector that carries the GFP reporter gene as well as the 4 Yamanaka genes.

Learning and spatial memory performance were assessed by means of the Barnes maze test. The learning performance of the OSKM-treated old rats was significantly improved compared to that of the control old counterparts. A marginal (P=0.06) improvement in the spatial memory was recorded in the treated versus control old rats. OSKM gene expression induced no pathological changes in the brain. The morphology and number of hippocampal cell populations like astrocytes and mature neurons did not show any changes with the treatment in the old rats as compared with the control old counterparts. The rat pan tissue DNAm age marker revealed that old OSKM gene-treated rats show a trend towards a decrease in epigenetic age. The Limma package was used to assess differential methylation by fitting linear models to the methylation data for specific group comparisons. Comparison of differential methylation between old treated and old control hippocampal DNA samples identified 671 differentially methylated CpGs probes (DPMs) in the DNA of OSKM-treated hippocampi (p<0.05). Assessment of the DPMs in old versus young controls revealed the presence of 1,279 hypomethylated CpGs near the promoter regions in young hippocampi (versus old controls) and 914 hypermethylated CpGs near the promoter in young hippocampi compared to old control hippocampi. We found a subset of 174 hypomethylated CpGs in the hippocampal DNA from old OSKM rats and young controls both compared with old control hippocampi. This means that in the hippocampal DNA there is a common set of CpGs which are hypermethylated during aging and are demethylated by the OSKM genes. This observation suggested that in these 174 CpGs the hypermethylation induced by aging is reversed by the demethylation effect of the OSKM genes on the same 174 CpGs. This observation can be interpreted as a rejuvenation effect of the OSKM genes of the old hippocampal methylome. Our results extend to the rat the evidence that viral vector-mediated delivery of the Yamanaka genes in the brain has strong regenerative effects without adverse side effects.

## INTRODUCTION

Aging is associated with a progressive increase in the incidence of neurodegenerative diseases, both in laboratory animals and humans. In rats, aging is accompanied by degenerative and/or atrophic processes in the cholinergic system of the forebrain, as well as morphological changes which parallel a reduction in spatial learning capacity (**1**). Likewise, there is solid evidence that, in this species, the decline in learning capacity and spatial reference memory that occurs with age is preceded by a multiplicity of structural, cellular, and molecular alterations at the hippocampal level (**2, 3**), many of which are comparable to those that occur during the aging of the human brain. At molecular level, gene expression studies in aging rodents have documented significant changes in hippocampal genes related to cholesterol synthesis, inflammation, transcription factors, neurogenesis and synaptic plasticity (**4-8**). In the hippocampus of female rats, it was reported that 210 genes are differentially expressed in senile as compared with young counterparts, most of them being downregulated (**9**).

There is clear evidence that the Yamanaka genes and other pluripotency genes have a high therapeutic potential in the senile central nervous system affected by different neurodegenerative diseases (for a review see **10).**

Recent results revealed that the Yamanaka genes display a dual behavior when expressed continuously *in vivo*, being regenerative when delivered via viral vectors but highly toxic when expressed in transgenic mice. Thus, it has been reported that delivery of the OSK genes by intravitreally injecting a regulatable adeno-associated viral vector type 2 (AAV2) expressing the polycistron OSK, can reverse vision deficits in an experimental model of glaucoma in mice as well as in 11 months old mice showing age-related vison impairment. Fifteen-months of continuous expression of OSK genes in retinal ganglion cells (RGC), induced neither pathological changes nor RGC proliferation. Intravenously OSK-AAV2-injected young and middle-aged mice (for 15 months) did not show any adverse side effects. In contrast, DOX-induced expression of the OSK genes in mice transgenic for OSK, induced rapid weight loss and death, likely due to severe dysplasia in the digestive system (**11**).

It has recently been reported that long-term gene therapy in the hypothalamus of young female rats, using an adenovector harboring the OSKM genes as well the GFP gene was able to attenuate the rate of reproductive decline so that the rate of fertility of 9 months old OSKM-treated females tripled the fertility of placebo vector treated counterparts (**12**). At 9 months of age female rats are near the age of ovulatory cessation (10 months). (**13**).

This line of evidence prompted us to implement medium-term (39 days) OSKM gene therapy in the dorsal hippocampus of old rats in order to restore learning and spatial memory performance in the experimental rats using as controls old counterparts injected with a placebo adenovector. The results of the study are reported here.

## MATERIAL AND METHODS

### Adenoviral Vectors

#### Construction of a Tet-Off regulatable HD-recombinant adenoviral vector (RAd) harboring the GFP and Yamanaka genes

The adenovector was constructed using a commercial kit (Microbix Inc., Ontario, Canada) that provides the shuttle plasmid pC4HSU, the helper adenovirus H14 and the HEK293 Cre4 cell line. A full description of the adenovector constructed has been already documented (**14**). Briefly, we cloned a construct harboring the bicistronic tandem Oct4-f2A-Klf4-ires-Sox2-p2A-cMyc (known as hSTEMCCA, generously provided by Dr. G. Mostoslavsky, Boston University) under the control of the bidirectional Tet-Off promoter PminCMV-TRE-PminCMV which on the second end is flanked by the gene for green fluorescent protein (GFP).

The hSTEMCCA cassette harbors the 4 Yamanaka genes grouped in pairs placed downstream and upstream of an internal ribosome entry site (IRES). In turn, each pair of genes is separated by a type 2A CHYSEL (cis-acting hydrolase element) self-processing short sequence which causes the ribosome to skip the Gly-Pro bond at the C-terminal end of the 2A sequence thus releasing the peptide upstream the 2A element but continuing with the translation of the downstream mRNA sequence. This allows near stoichiometric co-expression of the two cistrons flanking a 2A-type sequence (**15**). The whole expression cassette (STEMCCA cassette 10,065 bp) was cloned into the pC4HSU HD shuttle, giving rise to pC4HSU-STEMCCA-tTA. The pC4HSU HD shuttle consists of the inverted terminal repeats (ITRs) for Ad 5 virus, the packaging signal and part of the E4 adenoviral region plus a stuffer non-coding DNA of human origin which keeps a suitable size (28-31 Kbp) of the viral DNA.

The linearized DNA backbone of the new HD-RAd **(Fig 1)** was transfected in Cre 293 cells. For expansion, the helper H14 adenovirus was added to the cell cultures at a multiplicity of infection (MOI) of 5. In H14, the packaging signal is flanked by lox P sites recognized by the Cre recombinase expressed by the 293 Cre4 cells. Therefore, the helper virus provides in trans all of the viral products necessary for generation of the desired HD-RAd. The infected 293 Cre4 cells were kept until cytopathic effect (CPE) was evident. Cells were lysed and clear lysates mixed with H14 helper virus and added to a fresh culture of Cre4 293 cells at MOI 1. When CPE appeared, passage 2 (P2) cell lysates were prepared. This iterative co-infection process was carried on five more times in order to generate enough HD-RAd particles for virus purification. The newly generated HD-RAd was purified by ultracentrifugation in CsCl gradients. We used a control RAd harboring the hGFP gene.

**Figure 1.**
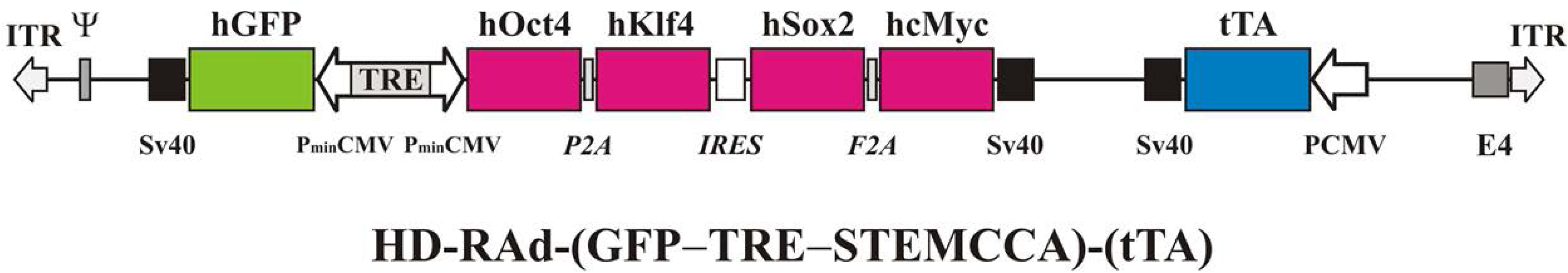
The figure illustrates the basic components of HD-RAd-STEMCCA-GFP-Tet-Off genome. Abbreviations-GFP: humanized Green Fluorescent Protein; TRE: Tetracycline responsive element; tTA: chimeric regulatory protein; PminCMV: cytomegalovirus minimal promoter; SV40pA: polyadenylation signal; ITR: inverted terminal repeats; ψ: packaging signal.

### Animals and in vivo procedures

Female Sprague-Dawley rats initially aged 3.5 (Y) and 25.3 (O) months were used. The animals were raised in our institution (INIBIOLP) and housed in a temperature-controlled room (22 ± 2°C) on a 12:12 h light/dark cycle (lights on from 7 to 19 h). Food and water were available ad libitum. All experiments with animals were performed according to the Animal Welfare Guidelines of NIH (INIBIOLP’s Animal Welfare Assurance No A5647-01).

#### Experimental design for long-term OSKM gene therapy in the dorsal hippocampus

Rats were allotted to a control or experimental group, thus forming 3 groups: Y intact (n=12), Old control (n=16) injected with RAd (GFP) and Old OSKM (n=17)-injected with RAd OSKM-GFP vector.

### Brain processing

Animals were placed under deep anesthesia and perfused with phosphate buffered para-formaldehyde 4%, (pH 7.4) fixative. The brains were rapidly removed and stored in para-formaldehyde 4%, (pH 7.4) overnight (4 °C). Finally, brains were maintained in cryopreservative solution at -20 °C until use. For immunohistochemical assessment, brains were cut coronally in 40 µm-thick sections with a vibratome (VT1000S; Leica Microsystems, Wetzlar, Germany).

### Immunohistochemistry

All immunohistochemical techniques were performed on free-floating sections. For each animal, separate sets of sections were immunohistochemically processed using anti-glial fibrillary acidic protein (GFAP) monoclonal antibody 1:500 (G3893, Sigma, Saint Louis, Missouri), anti-doublecortin (DCX) goat polyclonal antibody 1:250 (c-18, Santa Cruz Biotech., Dallas, Texas), mouse anti-hOct4 (1:10, BD Pharmingen, San Jose, CA), mouse anti-hSox2 (1:40, BD Pharmingen, San Jose, CA) and mouse anti-hc-Myc (1:50, BD Pharmingen, San Jose, CA).

For detection, the Vectastain® Universal ABC kit (1:500, PK-6100, Vector Labs., Inc., Burlingame, CA, USA) employing 3, 3-diamino benzidine-tretrahydro-chloride (DAB) as chromogen, was used. Briefly, after overnight incubation at 4°C with the primary antibody, sections were incubated with biotinylated horse anti-mouse antiserum (1:300, BA-2000,Vector Labs.) or horse anti-goat antiserum (1:300, BA-9500, Vector Labs), as appropriate, for 120 min, rinsed and incubated with avidin-biotin-peroxidase complex (ABC Kit) for 90 min and then incubated with DAB. Sections were counterstained with Nissl method (0.5% cresyl violet solution at 37°C for 10 minutes) to visualize anatomical landmarks and mounted with Vectamount (Vector Labs) to use for image analysis.

### Immunofluorescence

The Immunofluorescence technique was based on the following primary antibodies, Anti-Synaptophysin antibody [YE269] (ab32127), anti-GFAP monoclonal antibody 1:500 (G3893, Sigma see above) and anti NeuN-Mouse monoclonal antibody [1B7] (ABCAM Cat# ab104224) and mouse anti-KLF4 Polyclonal Antibody (Invitrogen Cat #PA5-27441). Once incubated with the primary antibodies (slides were incubated with either Goat anti Mouse Alexa Fluor™ 647 (Invitrogen Cat # A-21237) or Alexa Fluor 647 AffiniPure F(ab’)₂ Fragment goat Anti-Rabbit IgG (H+L) (Jackson Labs, 111-606-144), overnight at 4 °C in a dark chamber. Then, sections were rinsed threefold in PBS and counterstained for 15 min with DAPI. Control negative sections were prepared by omitting the primary antibody.

### Image Analysis of DCX cells

In each hippocampal block, one every six serial sections was selected in order to obtain a set of non-contiguous serial sections spanning the dorsal hippocampus (240 µm apart). The number of cells was assessed in the dorsal hippocampus which is located between coordinates-2.8 mm to -4.5mm from the bregma (**16**). Images were captured using an Olympus video camera (DP70. Tokyo, Japan) attached to an Olympus BX-51 microscope, using a 40X magnification objective. The total number of cells was estimated using a modified version of the optical dissector method (**17. 18, 19)**. Individual estimates of the total bilateral neuron number (N) for each region were calculated according to the following formula: N = RQΣ · 1/ ssf · 1/asf · 1/tsf, where RQΣ is the sum of counted neurons, ssf is the section sampling fraction, asf is the area sampling fraction, and tsf is the thickness sampling fraction.

Immunofluorescence images were captured using a laser scanning confocal microscope (Olympus FV1000, Japan). Neurons, neuropil and glial cells labeled with Alexa 647 were excited using a 635 nm solid state laser. All nuclei stained with DAPI were excited using a 405 nm solid state laser.

#### Stereotaxic injections

Rats were anesthetized with ketamine hydrochloride (40 mg/kg; ip) plus xylazine (8 mg/kg; im) and placed on a stereotaxic apparatus. In order to access the dentate gyrus (DG) of the dorsal hippocampus, the tip of a 26G needle fitted to a 10µl syringe was brought to the following coordinates relative to the bregma: AP: 3.5 mm ; ML: ± 2 mm; DV: 4 mm (**16**) and 3µl vector suspension per side was slowly injected (1µl/min).

### Spatial memory assessment

**Description of the Barnes maze protocol used-** The modified Barnes maze protocol used in this study was based on a previously reported procedure (**3, 18, 19**). It consists of an elevated (108 cm to the floor) black acrylic circular platform, 122 cm in diameter, containing twenty holes around the periphery. The holes are of uniform diameter (10 cm) and appearance, but only one hole is connected to a black escape box (tunnel). The escape box is 38.7 cm long x 12.1 cm wide x 14.2 cm in depth and it is removable. A white squared starting chamber (an opaque, 20 cm x 30 cm long, and 15 cm high, open-ended chamber) was used to place the rats on the platform. Four proximal visual cues were placed in the room, 50 cm away from the circular platform. The escape hole was numbered as hole 0 for graphical normalized representation purposes, the remaining holes being numbered 1 to 10 clockwise, and −1 to −9 counterclockwise **(Fig. 3-Inset)**. Hole 0 remains in a fixed position, relative to the cues in order to avoid randomization of the relative position of the escape box. A 90-dB white-noise generator and a white-light 500 W bulb provided the escape stimulus from the platform. We used an abbreviated protocol based on three days of acquisition trials (AT), making a total of 6 AT. An AT consists of placing a rat in the starting chamber for 30 s, the chamber is then raised, and the aversive stimuli (bright light and high pitch noise) are switched on and the rat is allowed to freely explore the maze for 120 s.

The behavioral performances were recorded using a computer-linked video camera mounted 110 cm above the platform. The video-recorded performances of the subjects were measured using the Kinovea v0.7.6 (http://www.kinovea.org) and Image Pro Plus v5.1 (Media Cybernetics Inc., Silver Spring, MD) software. The behavioral parameters assessed were as follows.

**(a) Escape box latency:** time (in s) spent by an animal since its release from the start chamber until it enters the escape box in an AT.
**(b) Goal sector (GS)**: Is the area of the platform corresponding to a given number of holes. GS3, is the area corresponding to holes -1, 0, +1.
**(c) Spatial memory assessment**: One day after the last AT is done, a trial, known as probe trial (PT), where the escape box has been removed, is performed, its purpose being to assess the permanence time in GS3, calculated taking the time spent by the rats in the area covered by the 3 holes during the PT.

### Hippocampus Dissection

Five rats from each group were sacrificed by decapitation, the brain was removed carefully, severing the optic nerves and pituitary stalk and placed on a cold plate. The hippocampus was dissected from cortex in both hemispheres using forceps. This dissection procedure was also performed on the anterior and posterior blocks, alternatively placing the brain caudal side up and rostral side up. After dissection, each hippocampus was immediately placed in a 1.5 ml tube and momentarily immersed in liquid nitrogen, then stored at -80 °C for DNA extraction.

### Hippocampal DNA extraction

Hippocampal DNA was extracted using an automated nucleic acid extraction platform called QIA cube HT (Qiagen) with a column-based extraction kit, QIA amp 96 DNA QIA cube HT Kit (Qiagen).

### Purification and methylation analysis of genomic DNA

DNAs from 24 hippocampi were purified using the DNeasy Blood and Tissue Kit (Qiagen). Only DNA samples with 260/280 ratios greater than 2.0 were processed for methylation analysis. The bisulfite conversion of genomic DNA was performed using the EZ Methylation Kit (Zymo Research, D5002), following the manufacturer’s instructions. The converted genomic DNA was analyzed by Illumina Infinium Horvath Mammal Methyl Chip40 at the UNGC Core Facility. The Horvath Mammal Methyl Chip40 assay provides quantitative measurements of DNA methylation for 22528 CpG dinucleotides that map to the Rattus norvegicus UCSC 6.0 genome (and other mammalian sequences) (**20**).

### Determination of epigenetic aging

In the present study, we used two rat clocks: rat brain clock and the rat pan-tissue, which were two of six rat clocks previously described (**21**). Briefly, the rat pan-tissue epigenetic clock was developed by regressing chronological age on CpGs from multiple rat tissues that are known to map to the genome of Rattus norvegicus. Age was not transformed. Penalized regression models were created with the R function “glmnet” (**22**). We also evaluated a mouse clock for brain samples which was described by Mozhui 2022 (23).

### Differential Methylation Analysis of Hippocampal DNA at Probe Level

In this study, probe-wise differential methylation analysis was performed using the Minfi and Limma packages in the R/Bioconductor environment. The Minfi package allowed for quality control, preprocessing, and analysis of DNA methylation data from Illumina microarrays. After preprocessing, the Limma package was used to assess differential methylation by fitting linear models to the methylation data for specific group comparisons: Old control versus Old OSKM hippocampi and Old control versus Young hippocampi. To ensure biological robustness, only CpG sites located within the proximal promoter region (<=1kb and 1-2kb upstream of the transcription start site) of relevant genes were considered. Additionally, to gain insights into the biological processes associated with the differentially methylated CpGs, gene ontology and pathway analysis using the Enrichr package in R/Biconductor were performed.

## RESULTS

### Effect of hippocampal OSKM gene therapy on spatial memory performance of old rats

Thirty days after the OSKM or control vector injection in the old rats, the 3 groups were submitted to the Barnes test in order to assess learning performance and spatial memory. As expected, learning performance **(Fig 2)** and spatial memory **(Fig 3)** declined with age. In the old rats, 39 days post OSKM gene therapy in the hippocampus induced a significant improvement in learning performance in the last two AT **(Fig 2. Right bar plots)**. In some of the old treated rats the videos recording their performance show a remarkable recovery. In the old rats hippocampal OSKM gene therapy induced a trend (P=0.06) towards an increase in spatial memory as compared with old controls **(Fig 3)**.

**Figure 2.**
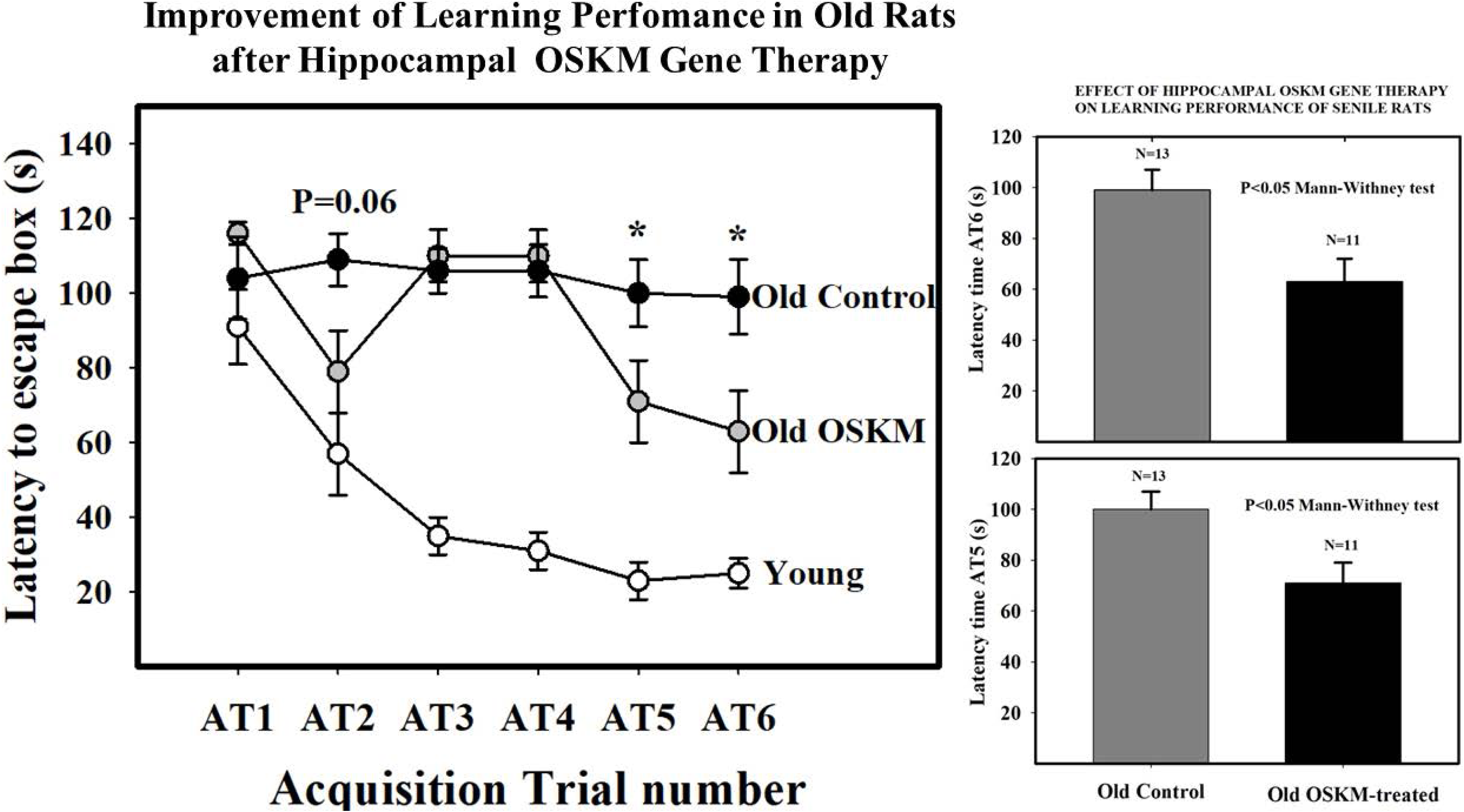
Effect of OSKM genes on learning performance in old rats. **Main plot** shows escape hole latency throughout training in young intact, old control and old OSKM treated rats. The training consists of six training sessions (acquisition trials, AT) in which the animals are given 120 seconds per AT to learn the location of the escape box, placed underneath hole 0. The latency time (the time, in s, that takes the animal to find the escape box since it is released from the starting box). Latency time falls rapidly in the young rats while in control old rats latency time remains as a plateau line during the whole training period. In contrast, old treated rats show little improvement from AT1 through AT4 but at AT5 and AT6 show a significant fall in latency time as compared with old controls. **Bar plots on the right** show the differences between old controls and old treated rats at AT5 and AT6.

**Figure 3.**
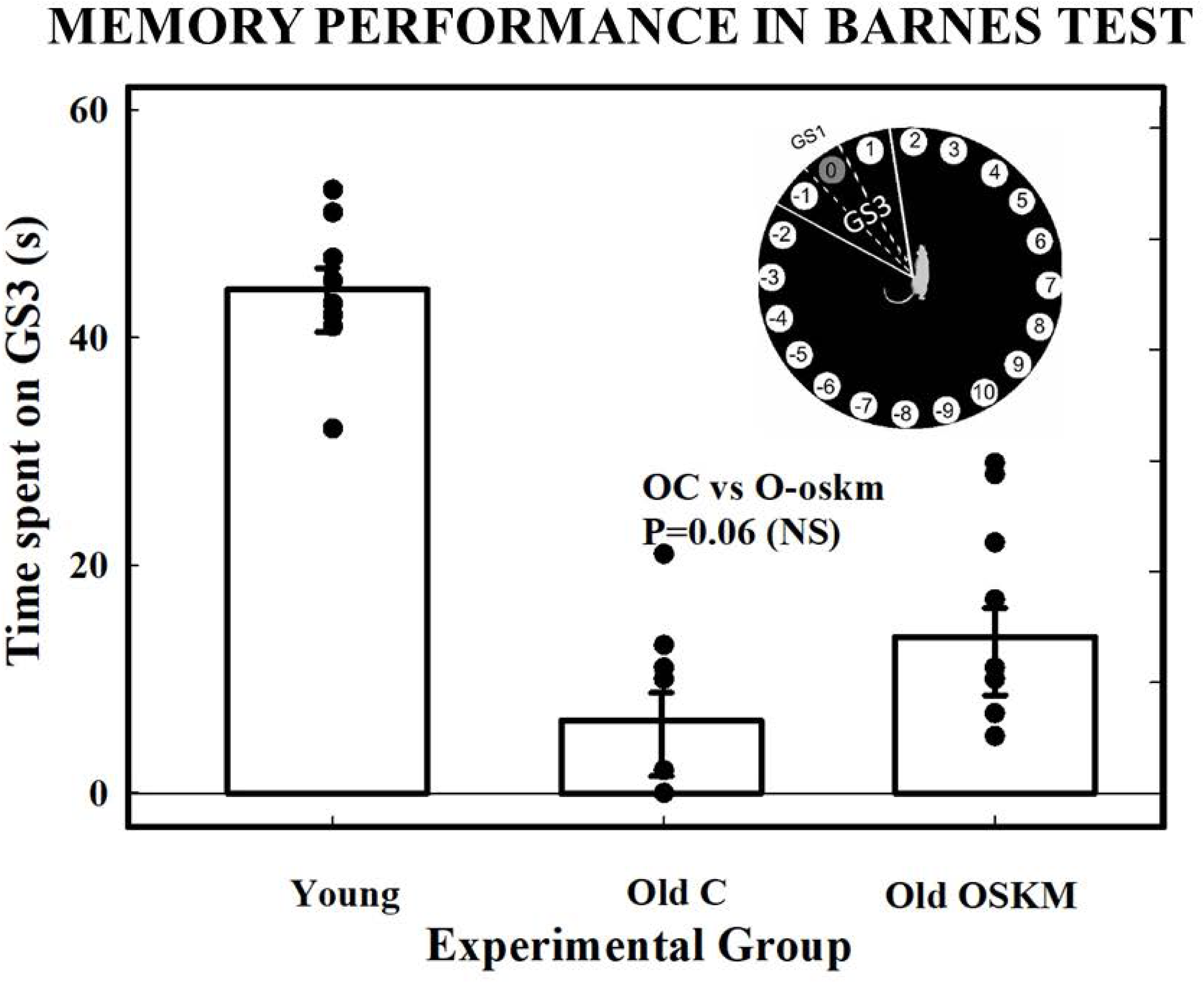
Spatial memory performance in the Barnes test. The main plot shows the time spent by rats in the GS3 sector exploring hole -1, 0 and +1 when the escape box is removed form hole 0. This is a reliable index of memory performance. While there is a marked fall in spatial memory between young and old rats, the OSKM treated old rats show a trend (P=0.06) towards an improvement versus old control animals. **The inset** shows a diagram of the Barnes platform delineating the GS3 sector.

### Effects of long-term OSKM gene expression on hippocampal tissue morphology and cell populations

In preliminary experiments it was confirmed that after OSKM vector injection GFP expression remains high for at least 30 days **(Fig. 4)**. OSKM gene expression was detected in the dentate gyrus (DG) for at least 3 weeks post vector injection **(Fig. 5).**

**Figure 4.**
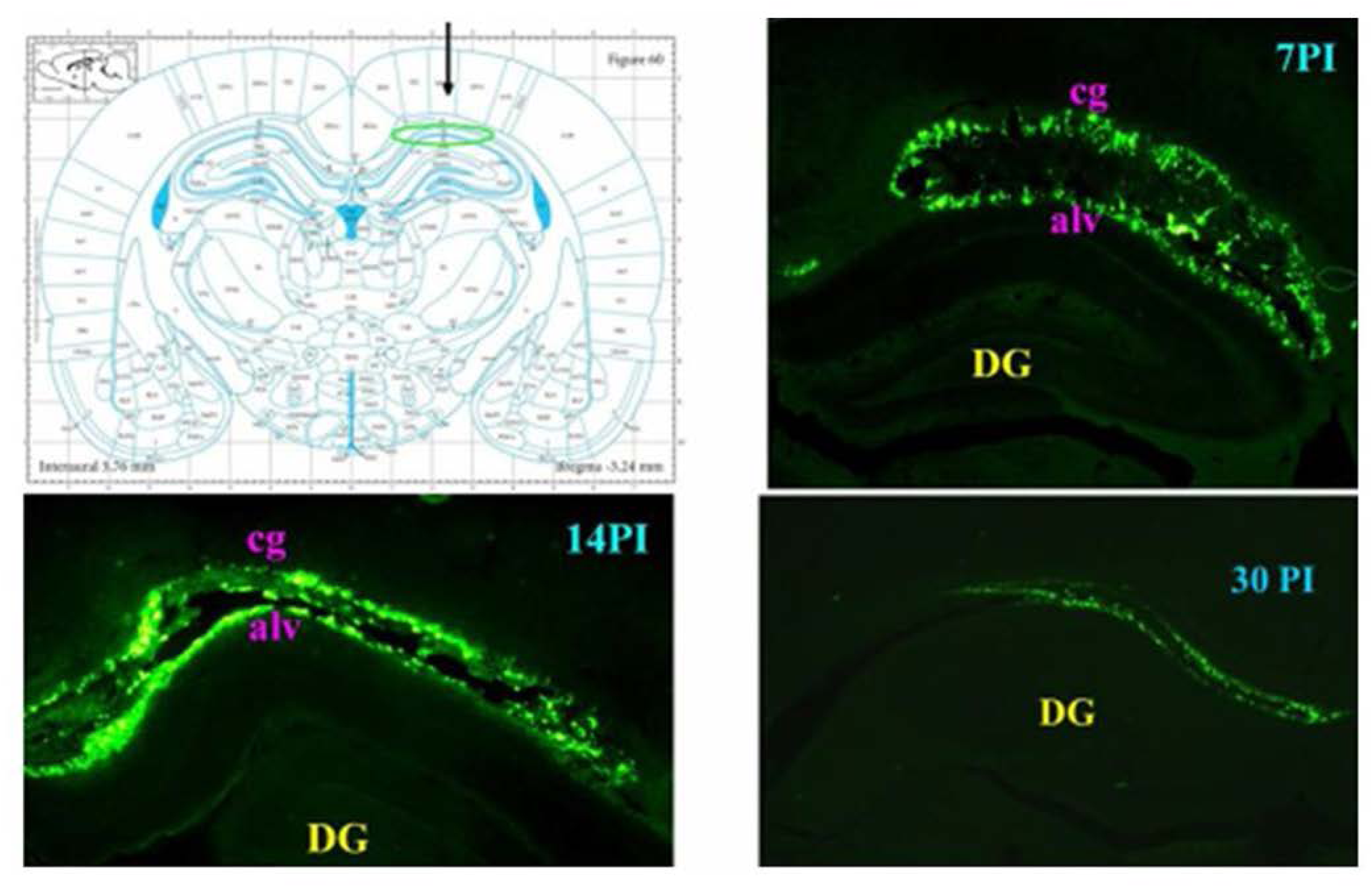
In the upper left diagram the arrow shows the region of the dentate gyrus of the hippocampus where the OSKM-GFP adenovector was delivered. The three additional panels show GFP fluorescence at different days post vector injection (PI), PI-7, PI-14 and PI-30.

**Figure 5.**
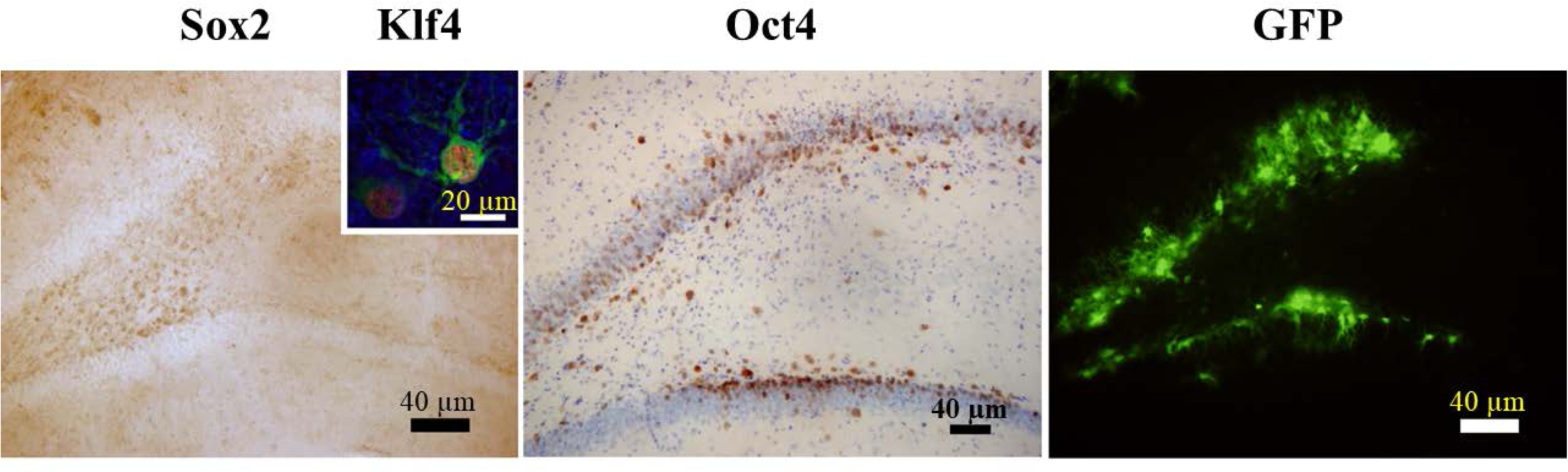
Expression of the Yamanaka genes in the hippocampus of OSKM-GFP adenovector-injected rats. The three main panels, from left to right, show expression of Sox2, Oct4 and GFP in the DG, 21 days after adenovector injection. **The inset** shows an hippocampal cell expressing Klf4 and GFP fluorescence. Scale bars show the corresponding magnification.

We were particularly interested in the possible effect of OSKM gene delivery to the granule cell layer of the DG. These migratory neurons were immunolabeled with the doublecortin (DCX) protein marker. DCX neurons are also found in the neurogenic subventricular zone (SVZ) of the lateral ventricle. Neurogenesis of DCX neurons is considered important for certain aspects of spatial memory. Their numbers decline markedly with age as illustrated by the bar plot shown on **Fig. 6**. In the old rats OSKM gene therapy did not affect the number of DCX neurons in the DG. In the main panels of **Fig. 6** it is also clear that 39 days of continuous OSKM expression did not induce any pathological changes in the hippocampal parenchyma (nor in any other region of the brain).

**Figure 6.**
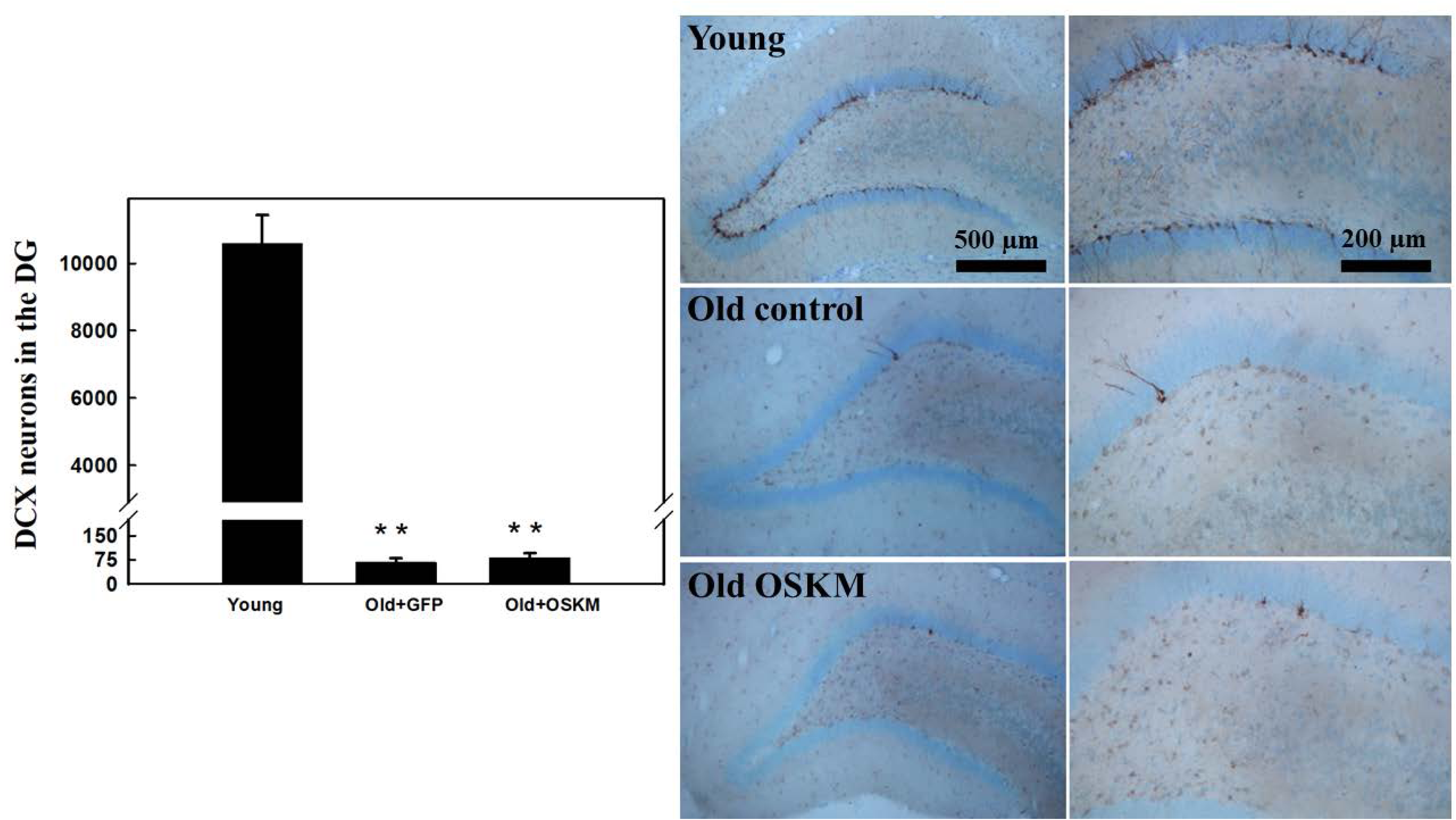
Effect of age and OSKM gene therapy on the granule cell layer of the DG of rats. The bar plot on the left shows the number of doublecortin (DCX) neurons in the DG of young, old control and Old OSKM treated rats. In old rats aging causes a remarkable fall in the number of DCX neurons. In contrast, the OSKM vector injection in the dorsal hippocampus of old rats has no significant effect on DCX cell number, 39 days post vector injection. The right panels illustrate the fall in DCX neuron number in old rats. It also shows that the hippocampal parenchyma of OSKM-treated old rats shows no pathological alterations due to OSKM expression for 39 days.

The morphology and number of other hippocampal cell populations like astrocytes (GAFP) and mature neurons (NeuN cells) did not show any changes with the treatment in the old rats as compared with the control old counterparts (**Fig 7**). Dentate gyrus synaptophysin-positive presynaptic bouton number was not affected either by the OSKM treatment in old rats (**Fig 7**, bottom panel).

**Figure 7.**
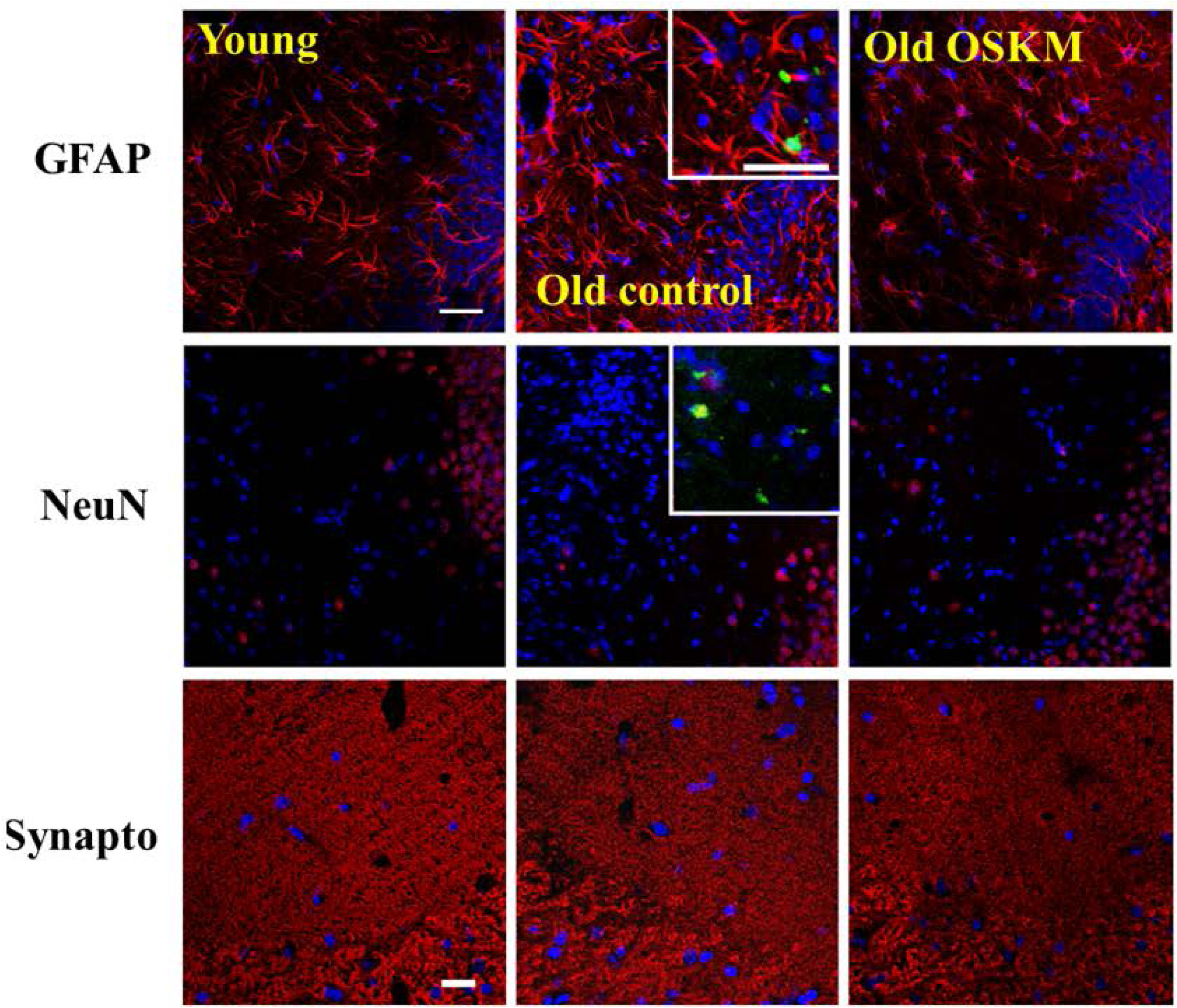
Effect of OSKM gene therapy on the hippocampal astrocyte and mature neuron population as well as on synaptic button density in old rats. After 39 days of OSKM vector injection in the dorsal hippocampus of old rats, there were no detectable changes in the astrocyte and mature neuron populations. Synaptophysin-positive presynaptic bouton density was not affected either. Insets show astrocytes or NeuN neurons expressing the GFP gene reporter.

### Epigenetic clock analysis of OSKM gene therapy

Epigenetic clocks are statistical models used to predict biological age based on patterns of methylation in cytosines. Our trio of epigenetic clocks were successful in differentiating between young and old hippocampal samples (Fig. 8, upper panel). However, when we limited our analysis to high-quality DNA samples from the hippocampi of older rats (with a sample size of 4 controls versus 5 rats treated with OSKM), the evidence for epigenetic rejuvenation based on our epigenetic clocks was merely suggestive. The rat brain clock showed a p-value of 0.08, and the mouse brain clock showed a two sided p-value of 0.057 (Fig. 8D,F). The pan-tissue clock, which was used for the rat samples, did not yield a significant difference, with a p-value of 0.17 (Fig. 8E).

**Figure 8.**
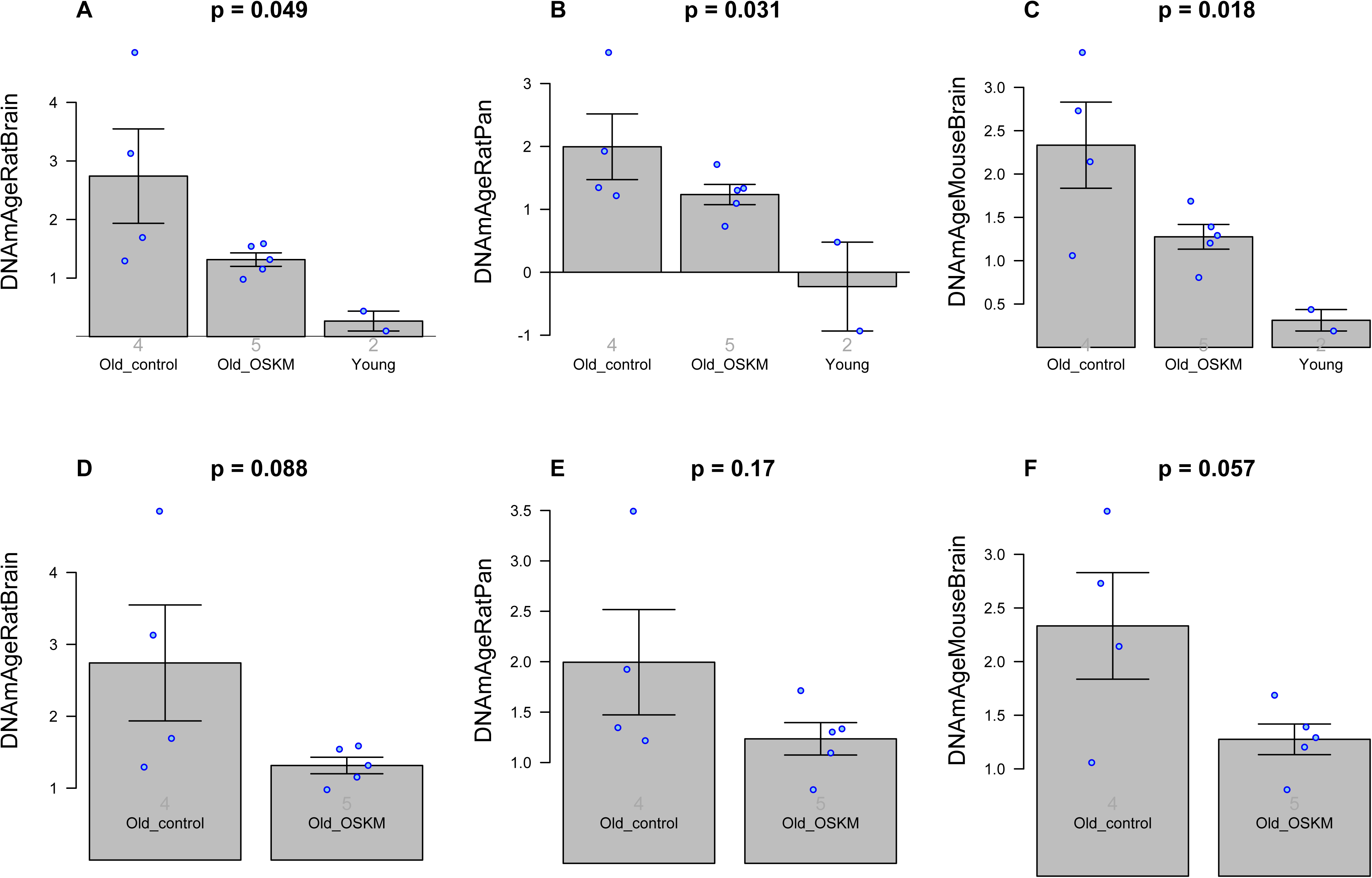
Epigenetic Clock analysis of OSKM gene therapy in the rat hippocampus. The columns correspond to three epigenetic clocks: A,D. Brain clock for rats. B,E. Pan tissue clock for rats. C,F. Brain clock for mice. All age estimates are in units of years (y-axis). A,B,C. report results for 3 conditions: young, old OSKM treated, and old control. D,E,F) The second row is analogous to the first row but the young group was removed. The title of each panel reports the nominal two sided Student test p-value. The faint grey numbers in each bar report the number of DNA samples per group (4 old controls, 5 old OSKM, 2 young). The bar plots report the mean value and one standard error. The rat clocks described in Horvath 2020 (**21**). The mouse clocks for brain samples is described in Mozhui 2022 (**35**).

**Figure 9:**
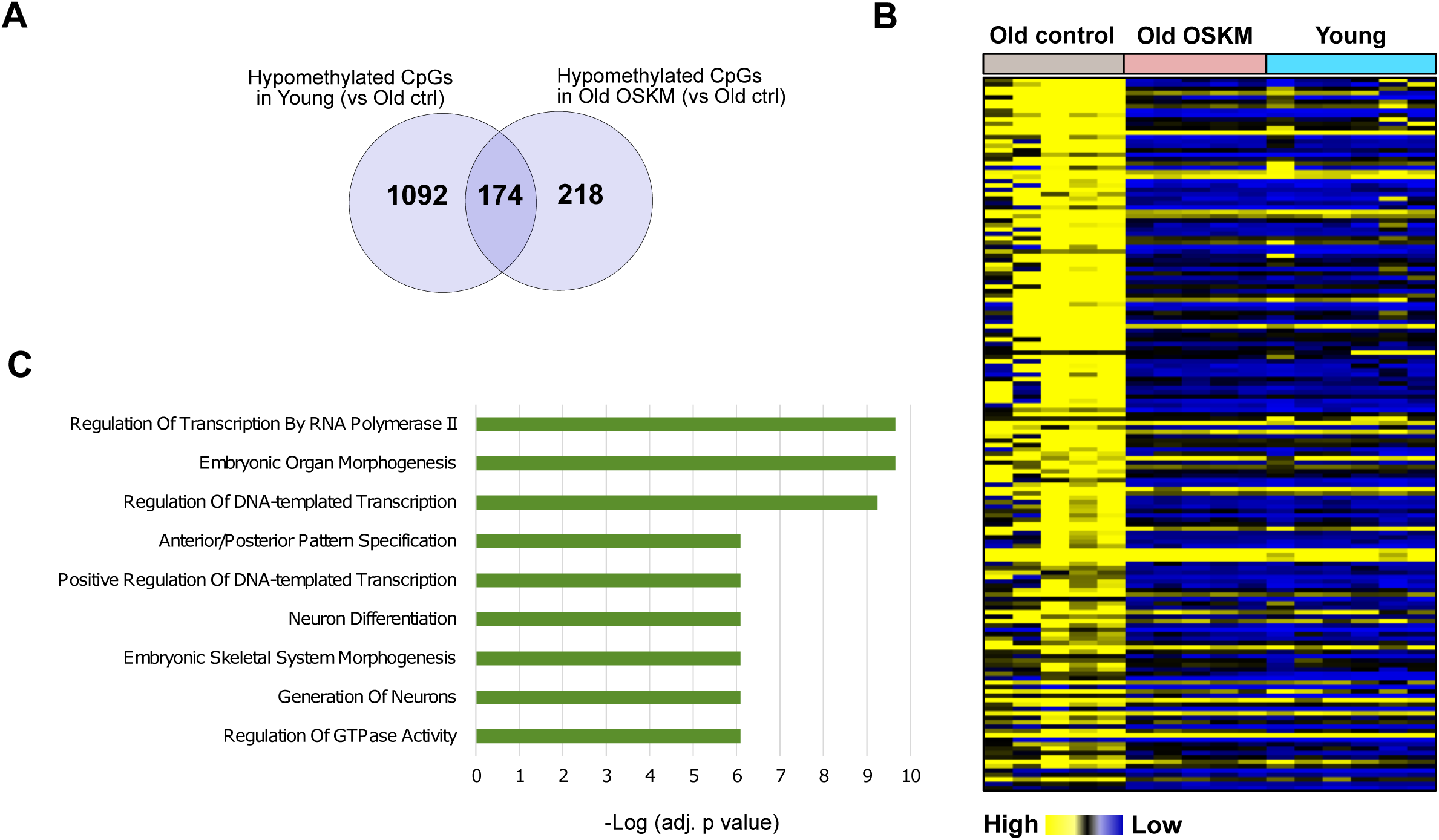
Common Hypomethylated CpGs in Young and OSKM-Treated Hypothalami Compared to Old Control Hypothalami. **A**. Venn Diagram Illustrating Differentially Hypomethylated CpGs in Young and OSKM-Treated Hippocampi Compared to Old Control Hippocampi **B**. Heatmap Representation of Beta Values for the 174 Commonly Hypomethylated Probes in Young and OSKM Groups Compared with Old Control. Visually, it is evident that for these 174 CpGs, the OSKM-treated and young hippocampi share a much closer methylation pattern and show marked differences with the methylation profile of the 174 CpGs of the Old control hippocampi. **C**. Barplot Illustrating Functional Enrichment of Genes Associated with the 174 CpGs

### Differential hippocampal DNA methylation in OSKM-treated old rats

Comparison of differential methylation between old treated and old control hippocampal DNA samples identified 671 differentially methylated CpGs probes (DPMs) in the DNA of OSKM-treated hippocampi (p<0.05; Supplementary file). Notably, the genes associated with hypomethylated CpGs near their promoter region were primarily involved in critical biological processes such as neuron differentiation and neuron development, indicating their potential role in shaping neuronal plasticity and cognitive function (Supplementary file 1). Some of them are NTRK2 - neurotrophic receptor tyrosine kinase 2, which is a gene that encodes a member of the neurotrophic tyrosine receptor kinase (NTRK) family. This kinase is a membrane-bound receptor that, upon neurotrophin binding, phosphorylates itself and members of the MAPK pathway.

NR4A2 - nuclear receptor subfamily 4 group A member 2, a gene that encodes a member of the steroid-thyroid hormone-retinoid receptor superfamily. The encoded protein may act as a transcription factor.

WNT1 - Wnt family member. The WNT gene family consists of structurally related genes which encode secreted signaling proteins. The gene is a member of the WNT gene family, highly conserved in evolution. The studies in mouse indicate that the Wnt1 protein functions in the induction of the mesencephalon and cerebellum.

POU3F2 - POU class 3 homeobox 2. This gene encodes a member of the POU-III class of neural transcription factors. The encoded protein is involved in neuronal differentiation and enhances the activation of corticotropin-releasing hormone regulated genes.

No processes were significantly found associated with hypermethylated CpG near their promoter in the OSKM-treated hippocampi (Supplementary file 1).

Assessment of the DPMs in old versus young controls revealed the presence of 1,279 hypomethylated CpGs near the promoter regions in young hippocampi (versus old controls) and 914 hypermethylated CpGs near the promoter in young hippocampi compared to old control hippocampi (p<0.05, Supplementary file 1). The hypomethylated genes in the young samples were also associated with processes related to the generation of neurons and neuron differentiation (Supplementary file 1). On the other hand, the hypermethylated genes in the young samples were linked to nervous and central nervous system development (Supplementary file 1).

Furthermore, we identified 174 CpG probes that exhibited common hypomethylation in both young and OSKM-treated old hippocampi when compared to old control hippocampi (**Supplementary file 1, Fig 9).** Functional enrichment analysis of the genes associated with these probes revealed processes related to neuron generation and differentiation, as well as embryonic morphogenesis, among others **(Supplementary file 1, Fig 9).** These findings provide valuable insights into the shared molecular mechanisms underlying neurodevelopmental processes in both young individuals and those subjected to OSKM treatment.

## DISCUSSION

There is documented evidence that in vivo partial cyclic reprogramming with OSKM genes in transgenic mice has regenerative effects. Thus, it was reported that the treatment applied to transgenic progeric mice, significantly prolonged their survival and partially rejuvenated some tissues although it did not rejuvenate the mice themselves (**24**). In another report, cyclic partial reprogramming in OSKM-transgenic, but otherwise wild type, middle-aged mice partially reversed the age-dependent reduction in hippocampal histone H3K9 trimethylation. The treatment elevated the levels of migrating granular cells in the DG and also significantly improved mouse performance in the object recognition test (**25**). Although the results of the latter study are in line with our present findings, our approach capitalized on the fact that Yamanaka gene therapy, unlike OSKM or OSK transgenic animals, allows continuous long-term expression of the genes, at least in the brain, without any adverse effects on cells or tissues, and exerts stronger and more prolonged regenerative effects.

In the present study we were interested in exploring the effects of OSKM gene delivery on hippocampal DNA methylation. We have discovered suggestive evidence of epigenetic rejuvenation, as indicated by two distinct epigenetic clocks specifically designed for rat and mouse brain samples. This evidence aligns with a multitude of other studies that have similarly reported signs of epigenetic rejuvenation in human and mouse cells and tissues after the application of OSKM or OSK. (**26-31**)

Our investigation using epigenetic clocks was somewhat constrained by the limited sample size, including 4 controls compared to 5 OSKM-treated samples. Nonetheless, these epigenetic clocks, which have been trained using independent data, were previously evaluated and stand out in terms of their accuracy Early studies performed in rat brain had shown a global loss of DNA methylation during aging (**32**) but there was limited information on the DNAm landscape in the rat hippocampus and even less about the impact of aging on this landscape. In a recent study in young (2.6 mo) and old (26.6 mo) rats we found that 1,090 CpGs exhibited increased methylation in the hippocampal DNA of the old rats. Additionally, an enrichment pathway analysis revealed that neuron fate commitment, brain development, and central nervous system development were processes whose underlying genes were enriched in hypermethylated CpGs in the old animals (**33**). In the old rat hippocampi of the cited study, the methylation levels of CpGs proximal to transcription factors associated with genes Pax5, Lbx1, Nr2f2, Hnf1b, Zic1, Zic4, Hoxd9; Hoxd10, Gli3, Gsx1 and Lmx1b, and Nipbl showed a significant inverse regression with spatial memory performance. Furthermore, regression analysis of different memory performance indices with hippocampal DNAm age was significant. Those results suggested that age-related hypermethylation of transcription factors related to certain gene families, like Zic and Gli, may play a causal role in the decline in spatial memory in old rats.

Although in the present study we did not find evidence that expression of the OSKM genes in the hippocampus of old rats modified the methylation levels of memory-related genes families like Zic or Gli, we did find a subset of 174 hypomethylated CpGs in the hippocampal DNA from old OSKM rats and young controls both compared with old control hippocampi. This means that in the hippocampal DNA there is a common set of CpGs which are hypermethylated during aging and are demethylated by the OSKM genes. This observation can be interpreted by stating that in these 174 CpGs the hypermethylation induced by aging is reversed by the demethylation effect of the OSKM genes on the same 174 CpGs. The observation is consistent with the well-established rejuvenation effects of OSKM genes. This hypomethylation effect of OSKM genes is likely to derepress a number of hippocampal genes some of which may be involved in the improvement of the learning performance and spatial memory documented by the results of the Barnes test. The results are also consistent with a report indicating that in the hippocampus of female rats 210 genes are differentially expressed in senile as compared with young counterparts, most of them being downregulated (**9**). There is virtually no information about the impact of OSKM (or OSK) gene expression in the hippocampus of old animals (or humans) on the DNA methylation landscape. Our results now show that, while normal hippocampal aging prevalently hyper methylate some genes, OSKM gene therapy hypomethylates a sub set of genes which could lead to increased expression of hippocampal genes that favor cognitive performance in old rats. Clearly, available experimental data are insufficient to elucidate the mechanism by which OSKM genes exert their beneficial effect on cognitive performance.

Concerning the marginal epigenetic rejuvenation observed in the treated old hippocampi, it should be pointed out that only a small proportion of hippocampal cells are expected to have been transduced by our OSKM-GFP vector and therefore directly rejuvenated. In addition, those few cells expressing the OSKM genes may have generated, via the release of revitalizing factors, a rejuvenating environment in the hippocampal tissue, contributing to mildly reduce the epigenetic age of the whole region.

## CONCLUDING REMARKS

Our results extend to the rat the evidence that viral vector-mediated delivery of the Yamanaka genes in the brain has strong regenerative effects without adverse side effects. Our HD OSKM-GFP adenovector constitutes a suitable gene delivery tool for the long-term expression of large polycistronic gene constructs. Using the same adenovector, we have recently shown that 5.8-month long expression of OSKM in the hypothalamus of young female rats attenuates the age-related decline in fertility at 9 months of age (rats stop ovulating at 10 months) an achievement not previously documented (**12**).

Since one of the evolutionary purposes of the Yamanaka genes is rejuvenation (34,35) experimental studies aimed at exploring their functional effects in adult and old animals seem a promising avenue of research in regenerative medicine.

